# Accelerated Aging after Traumatic Brain Injury: an ENIGMA Multi-Cohort Mega-Analysis

**DOI:** 10.1101/2023.10.16.562638

**Authors:** Emily L Dennis, Samantha Vervoordt, Maheen M Adamson, Houshang Amiri, Erin D Bigler, Karen Caeyenberghs, James H Cole, Kristen Dams-O’Connor, Evelyn M Deutscher, Ekaterina Dobryakova, Helen M Genova, Jordan H Grafman, Asta K Håberg, Torgeir Hollstrøm, Andrei Irimia, Vassilis E Koliatsos, Hannah M Lindsey, Abigail Livny, David K Menon, Tricia L Merkley, Abdalla Z Mohamed, Stefania Mondello, Martin M Monti, Virginia FJ Newcome, Mary R Newsome, Jennie Ponsford, Amanda Rabinowitz, Hanne Smevik, Gershon Spitz, Umesh M Vankatesan, Lars T Westlye, Ross Zafonte, Paul M Thompson, Elisabeth A Wilde, Alexander Olsen, Frank G Hillary

## Abstract

**Objective:** The long-term consequences of traumatic brain injury (TBI) on brain structure remain uncertain. In light of current evidence that even a single significant brain injury event increases the risk of dementia, brain-age estimation could provide a novel and efficient indexing of the long-term consequences of TBI. Brain-age procedures use predictive modeling to calculate brain-age scores for an individual using MRI data. Complicated mild, moderate and severe TBI (cmsTBI) is associated with a higher predicted (brain) age difference (PAD), but the progression of PAD over time remains unclear. Here we sought to examine whether PAD increases as a function of time since injury (TSI).

**Methods:** As part of the ENIGMA Adult Moderate and Severe (AMS)-TBI working group, we examine the largest TBI sample to date (n=343), along with controls, for a total sample size of 540, to reproduce and extend prior findings in the study of TBI brain age. T1w-MRI data were aggregated across 7 cohorts and brain age was established using a similar brain age algorithm to prior work in TBI.

**Results:** Findings show that PAD widens with longer TSI, and there was evidence for differences between sexes in PAD, with men showing more advanced brain age. We did not find evidence supporting a link between PAD and cognitive performance.

**Interpretation:** This work provides evidence that changes in brain structure after cmsTBI are dynamic, with an initial period of change, followed by relative stability, eventually leading to further changes in the decades after a single cmsTBI.

## Introduction

Complicated mild (mild TBI with trauma-related intracranial pathology on CT/MRI), moderate, and severe TBI (cmsTBI) results in downstream consequences for brain structure and physiology, altering the course of brain aging and increasing risk for neurodegeneration.^1,2^ The trajectories for brain atrophy in remote cmsTBI (>10 years) samples have not been studied extensively but some evidence suggests a pattern distinct from that in Alzheimer’s disease.^3^ Modifiers of aging after cmsTBI include chronic neuroinflammation,^4^ blood-brain barrier disruption, and proteinopathy primarily involving tau, beta-amyloid, and alpha-synuclein.^5,6^ As one ages with cmsTBI, the initial injury characteristics and time-since-injury (TSI) may therefore interact to moderate long-term outcomes.^7^

Brain-age-gap has been developed as a potential biomarker for outcome in psychiatric and neurological disorders. Brain-age is established by comparing brain characteristics of an individual to their chronological age, determined using data from healthy participants.^8,9^ Studies of the predicted age difference (PAD) have been applied to a range of clinical populations, including depression,^10^ PTSD,^11^ and stroke.^12^ Brain age prediction modeling shows that, within the first several years after cmsTBI, there is increased atrophy in gray and white matter equivalent to about half a decade in chronological age in msTBI,^13–16^ and 1-3 years in mild TBI.^2,17,18^ The extent to which post-traumatic atrophy may evolve as individuals transition through various stages of life, continuing into senescence for years and even decades, remains to be established.

Greater PAD after cmsTBI has been associated with greater injury severity and poorer cognitive function. However, it remains unclear whether the *brain atrophy* – observed in the well-documented sulcal and ventricular enlargement occurring over the first few years secondary to lesion resolution and transsynaptic or Wallerian degeneration^19^ – has long-term effects and is associated with an *acceleration* of brain aging over the lifespan. While there is some evidence of an association between TSI and PAD in mid-life, interpreted as *accelerated aging*,^13^ other work has failed to substantiate this effect.^15^

This study extends prior work^13–15^ by analyzing a larger sample with a wider age range (18-85) and TSI (mean=5.3 years ±6.4, range 0.2-34), with severity ranging between complicated mild and severe. We leveraged a team science approach through ENIGMA (Enhancing NeuroImaging Genetics through Meta-Analysis^20^) and the ENIGMA Adult msTBI (AMS-TBI) working group.^21^ We hypothesized that we would see both *advanced brain age* in individuals with cmsTBI, and *accelerated brain aging*, defined as increasing PAD as a function of TSI. We hypothesized that injury severity, lower educational attainment, and poorer cognitive function would be independent predictors of greater PAD. Additionally, we examined the influence of sex on brain age trajectory without making specific predictions, due to mixed findings in the literature.^22^ Finally, given its role as a potential risk factor for poor outcome after TBI,^23^ we examined the role of genetic risk (APOE) on PAD as it interacts with TSI, anticipating that the ε4 allele would confer risk for more accelerated brain aging.

## Methods

### Study samples

Study samples consisted of seven cohorts from parent studies originating in three countries (see **Table 1**). The final sample size (as detailed below and in **Figure 1**) was 540: 343 cmsTBI (237M/106F, mean age=44.5±16.2 years, range=20-85) and 197 control (113M/84F, mean age=38.6±16.2 years, range=18-84). There was a significant difference in age between groups (*p*<0.001), partially attributable to two cohorts with older participants not including a control sample. Participants were recruited from hospitals, outpatient rehabilitation clinics, and the surrounding community. Details on the inclusion and exclusion criteria for each cohort are included in **Supplementary Table 1**. Original studies were reviewed by the appropriate institutional review board for each respective institution. All participants provided written or verbal informed consent as part of involvement with the parent study.

**Figure 1.**
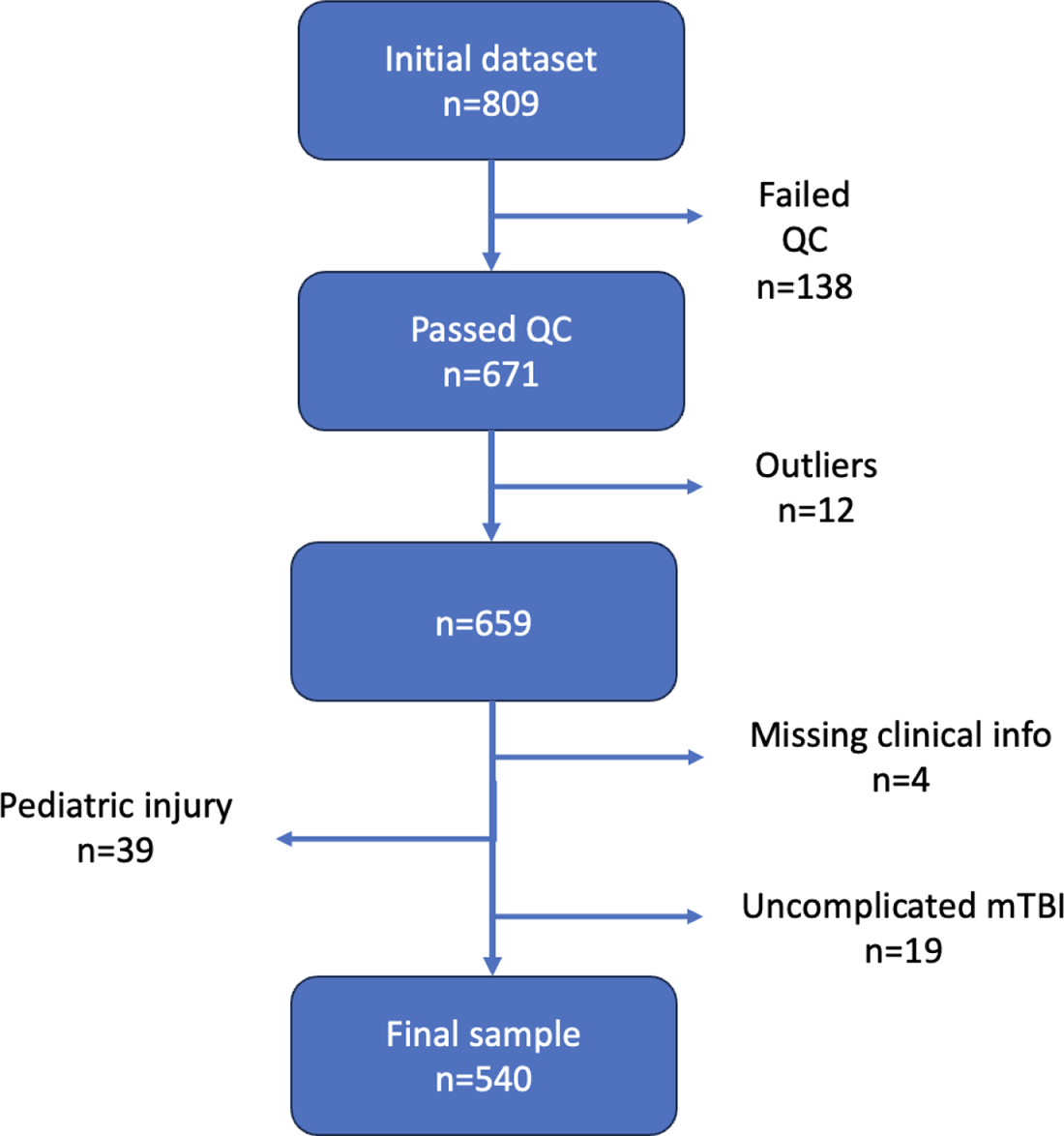
Participant exclusion flowchart. The number of participants included and various reasons for exclusion. This information is also detailed by site in **Supplementary Table 1.**

**Table 1.** Cohort Demographics. The total sample size, number of cmsTBI and control participants, male and female participants, average age (and standard deviation), average time since injury (TSI; in years, and standard deviation), and range of TSI are shown for each cohort.

| Cohort | Total N | Subgroup N |  | M/F | Age (SD) \| Range | TSI (in years; average \| range) |
| --- | --- | --- | --- | --- | --- | --- |
| Kessler | 63 | cmsTBI | 36 | 25/11 | 40.2 (11.2) \| 21-65 | 7.4 (5.8) \| 1.4-27 |
|  |  | Control | 27 | 12/15 | 41.2 (10.8) \| 21-63 | - |
| LETBI | 53 | cmsTBI | 53 | 30/23 | 58.0 (10.3) \| 40-85 | 11.1 (9.2) \| 1.2-34.4 |
|  |  | Control | 0 | - | - | - |
| Monash | 97 | cmsTBI | 74 | 60/14 | 38.3 (14.5) \| 20-78 | 2.1 (3.1) \| 0.2-21.1 |
|  |  | Control | 23 | 10/13 | 30.4 \| (12.6) 18-69 | - |
| NTNU | 113 | cmsTBI | 45 | 33/12 | 32.3 (11.3) \| 20-65 | 2.8 (1.1) \| 1.4-5.4 |
|  |  | Control | 68 | 50/18 | 35.2 (14.3) \| 19-64 | - |
| Oslo | 64 | cmsTBI | 64 | 43/21 | 42.9 (13.1) \| 20-67 | 1.2 (0.4) \| 0.6-1.8 |
|  |  | Control | 0 | - | - | - |
| PSU | 112 | cmsTBI | 53 | 36/17 | 55.4 (16.9) \| 20-79 | 9.3 (6.7) \| 0.6-23.6 |
|  |  | Control | 59 | 31/28 | 45.3 (20.2) \| 18-84 | - |
| VA Palo Alto | 38 | cmsTBI | 18 | 10/8 | 39.9 (13.6) \| 24-71 | 8.1 (5.5) \| 0.7-20.2 |
|  |  | Control | 20 | 10/10 | 37.4 (10.2) \| 23-54 | - |
| **Total** | **540** | **cmsTBI** | **343** | **237/106** | **44.5 (16.2) \| 20-85** | **5.3 (6.4) \| 0.2-34.4** |
|  |  | **Control** | **197** | **113/84** | **38.6 (16.2) \| 18-84** | **-** |

For the analyses in this study, only participants 18 years of age or older at the time of enrollment were included and participants who sustained their injury before the age of 18 were excluded from most analyses (N=61) in order to avoid confounding brain changes associated with neurodevelopment with morphological alterations attributed to injury. Level of education was measured using ISCED 2011 categories.^24^

Patients had to have sustained a cmsTBI, defined by having a TBI and trauma-related intracranial pathology and/or significant loss-of-consciousness (for details see below). Complicated mild TBI has been shown to result in more severe and chronic cognitive deficits compared to mild TBI/concussion, constituting a distinct class of injury,^25^ and was therefore included in the analysis. As injury severity was operationalized differently across the parent studies, patients were reclassified into complicated mild, moderate, or severe TBI based on GCS (Glasgow Coma Scale) score, where available: (1) GCS 14-15 and trauma-related intracranial pathology=complicated mild TBI, (2) GCS 9-13=moderate TBI, and (3) GCS 3-8=severe TBI. Where GCS scores were not available, injury severity was determined by other available study specific procedures (see **Supplementary Table 1**) or inferred based on inclusion/exclusion criteria. Severity was coded as 1=complicated mild TBI, 2=moderate TBI, 3=severe TBI.

### Image acquisition

All cohorts shared their raw T1-weighted MRI data with the central processing site (University of Utah, ED). The acquisition parameters for each cohort are shown in **Supplementary Table 2**.

### Cognitive data

Most sites collected a version of the Trail Making Test (TMT) and Digit Span from their cohorts, therefore we examined whether TMT condition A and B (or D-KEFS conditions 3 and 4) performance or Digit Span scores (forward + backward) was associated with PAD in the cmsTBI group. For TMT, using raw or scaled scores together was not appropriate given differences in Halstead Trails and D-KEFS Trails test administration and norming procedures. For this reason, we normed the data based on our healthy control subjects to calculate *T*-scores, separately for each test.

### Brain age prediction

We implemented the Gaussian processes regression approach for brain age estimation.^13^ Raw T1-weighted MR images were processed with the brainageR v2.1 workflow (https://github.com/james-cole/brainageR), with a model similar to those described previously.^11,13^ The brainageR model was trained on brain MRIs from 3,377 individuals from seven publicly available datasets. Overall, this training sample included individuals 18-92 years old from samples in the United States, United Kingdom, Australia, and China. Briefly, T1-weighted images were segmented into gray matter, white matter, and cerebrospinal fluid using SPM12 (https://www.fil.ion.ucl.ac.uk/spm/software/spm12/) and spatially normalized. The resulting images were vectorized and subjected to principal components analysis (using R *prcomp* https://cran.r-project.org), where components explaining the top 80% of variance were retained, resulting in 435 components. Processing for the ENIGMA AMS-TBI data was the same, with raw T1-weighted images segmented, normalized, vectorized, and the rotation matrix from the training dataset applied to yield 435 components for each participant. The resulting components were used to predict brain age using *kernlab*,^26^ and tissue segmentations were visually checked for quality. Of the 809 scans across seven cohorts, 17% failed visual quality control due to poor tissue segmentation (QC, N=138). The failure rate was similar between the cmsTBI and control groups. Outliers based on predicted age difference (PAD; ±3SD) were removed (N=12). A flowchart of reasons for exclusions may be seen in **Figure 1** and site-level information in **Supplementary Table 3**. Due to missing demographic or clinical information for some participants, the final sample size was 540 participants (343 cmsTBI and 197 controls).

The variable of interest was PAD, calculated by subtracting the chronological age from the predicted age. A negative PAD score indicates brain age values that are lower (younger) than expected given an individual’s chronological age. A positive PAD indicates a brain that appears older than expected, and could imply either advanced and/or accelerated aging. Plots of the chronological age and predicted brain age across cohorts are shown in **Supplementary** Figure 1. For the purposes of this paper, *advanced* brain aging refers to a larger PAD in TBI, while *accelerated* brain aging refers to a PAD that increases with more advanced chronological age or more time post-injury.

### APOE analyses

A total of 166 participants (128 cmsTBI) across 3 cohorts had available APOE genotype. Of 166 participants with APOE genotype, 56 had at least one ε4 allele. Due to the limited APOE sample size, all findings related to APOE were considered exploratory and should be interpreted with caution.

### Data availability

Data are available to researchers who join the working group and submit a secondary analysis proposal to the group for approval, which is granted on a cohort-by-cohort level. Interested researchers should contact the corresponding authors.

#### Statistical Analyses

Statistical analyses were run as mixed effects models in R 3.1.3 with the *nlme* package, setting PAD as the dependent variable. Some cohorts consisted of participants from multiple studies or sites. Nested random effects (intercepts) were used to control for cohort and site/study. A flowchart for the statistical models tested may be found in **Supplementary** Figure 2. Normalized residuals, accounting for age, sex, and random effects of cohort and site, were calculated from regression analyses and used for charting.

### Demographic variables

As a first test of model accuracy, we examined the correlation between predicted brain age and chronological age in the healthy control sample. The model accurately predicted chronological age in healthy individuals (r=0.92). Across the whole sample, a significant correlation between age and PAD was found (*r*=-0.2, *p*<.001), meaning that PAD was higher for younger participants. The negative association between age and PAD has been shown in numerous papers and may result when there is insufficient information to estimate brain age while attempting to minimize residuals, resulting in regression to the mean/median.^27^ There was not a significant sex difference in PAD (*t*(538)=-0.4, *p*=.66). Both age and sex were included as covariates in the models. Lastly, we examined the association between PAD and years of education, which was non-significant (*t*(526)=-1.9, *p*=.06).

### Primary group comparison

We examined differences in PAD between the cmsTBI and control group, covarying for chronological age and sex. We also compared controls to TBI groups with patients broken into severity categories - cmTBI, modTBI, and sevTBI.

### Sensitivity analyses

Several additional sensitivity analyses were run. First, we excluded cmsTBI participants with lesions visible on the T1-weighted image (n=152) and those who were scanned <1 year post-injury (n=51). The rationale for the former is that lesions could lead to errors in the processing pipeline, and we wanted to ensure results were not due to such bias. The rationale for the latter is that this is a dynamic period during which most recovery occurs, and there may be diaschisis-related atrophy that is distinct from the more long-term interaction between aging and TSI.^28^ We also ran analyses excluding individuals over 60 (age at scan, n=94) to check a sample with better age matching between patients and controls.

### Within-TBI-group analyses

We examined associations between PAD and a number of demographic, clinical, and cognitive variables within the cmsTBI group.

#### TSI

To examine a potential accelerated aging effect using cross-sectional data, we examined whether PAD remained consistent over TSI. We conceptualize accelerated aging as changing (advancing PAD) with increasing years since the time of injury. We tested linear and nonlinear association with PAD across all cmsTBI participants with TSI>1 year, TSI>5 years, and TSI>10 years. To determine the shape of nonlinear associations, we performed spline interpolation with the *gam* function in the *splines* R library, testing 3, 5, and 7 degrees of freedom. We tested this association in these distinct windows of TSI as we would expect that detectable evidence of accelerated aging may not emerge for a few years after injury. *That is, if the pathological consequences of TBI (proteinopathy, inflammation) are active in chronic TBI, these interacting and perhaps cumulative effects should be more evident with longer windows of TSI*.

#### Age

Chronological age and TSI are related, making it difficult to tease apart the individual effects of each on PAD, especially given the known association between chronological age and PAD. In addition, injury severity and age-at-injury are related (**Supplementary** Figure 3). To address this, we examined associations with TSI within separate age brackets, and associations with age within different TSI brackets.

#### Cognitive performance

We examined associations with cognitive function, specifically performance on the TMT task (measured using either Delis-Kaplan Executive Function System [D-KEFS] or Halstead-Reitan Neuropsychological Battery [HRNB]; harmonization described above) and Digit Span. There were 175 participants in the cmsTBI group with Digit Span data (Digit Span Forward and Backward - DSF and DSB) and 203 with TMT (81 with D-KEFS and 122 with HRNB, normed separately using 95 controls with D-KEFS and 72 with HRNB). These analyses were adjusted for age and sex.

### Interactions

We tested the following interactions: group × age, group × sex, and within the cmsTBI group, age × TSI, TSI × sex, and TSI × education.

### APOE analyses

We compared PAD between participants negative for ε4 alleles and individuals either homo- or heterozygous for the allele across the whole sample and within the cmsTBI group only.

### Defining survivor bias

Survivor bias is a pervasive methodological issue that faces any research efforts examining disease and mortality. This can result in research recruitment of an artificially healthy sample with respect to brain and behavioral health. Given the link between education and health, longevity, and mortality,^29,30^ we examined higher education in our older cohorts as a proxy for survivor bias.

## Results

### Primary group comparison

Across 540 individuals, the cmsTBI group had a significantly larger PAD than the control group (*b*=4.99, *p*<.001, **Figure 2a**), indicating a substantial deviation (5 years) between brain age and chronological age with older appearing brains (*advanced aging*) in cmsTBI. There was a significant association between PAD and injury severity group, with more severe injury being associated with greater PAD (*b*=1.51, *p*=.019, **Figure 2b**). Separated by severity, the cmTBI group showed the smallest group difference (cmTBI: N=288, *b*=2.3, *p*=.055; modTBI: N=248, *b*=4.4, *p*=<.001; sevTBI: N=304, *b*=5.8, *p*=<.001).

**Figure 2.**
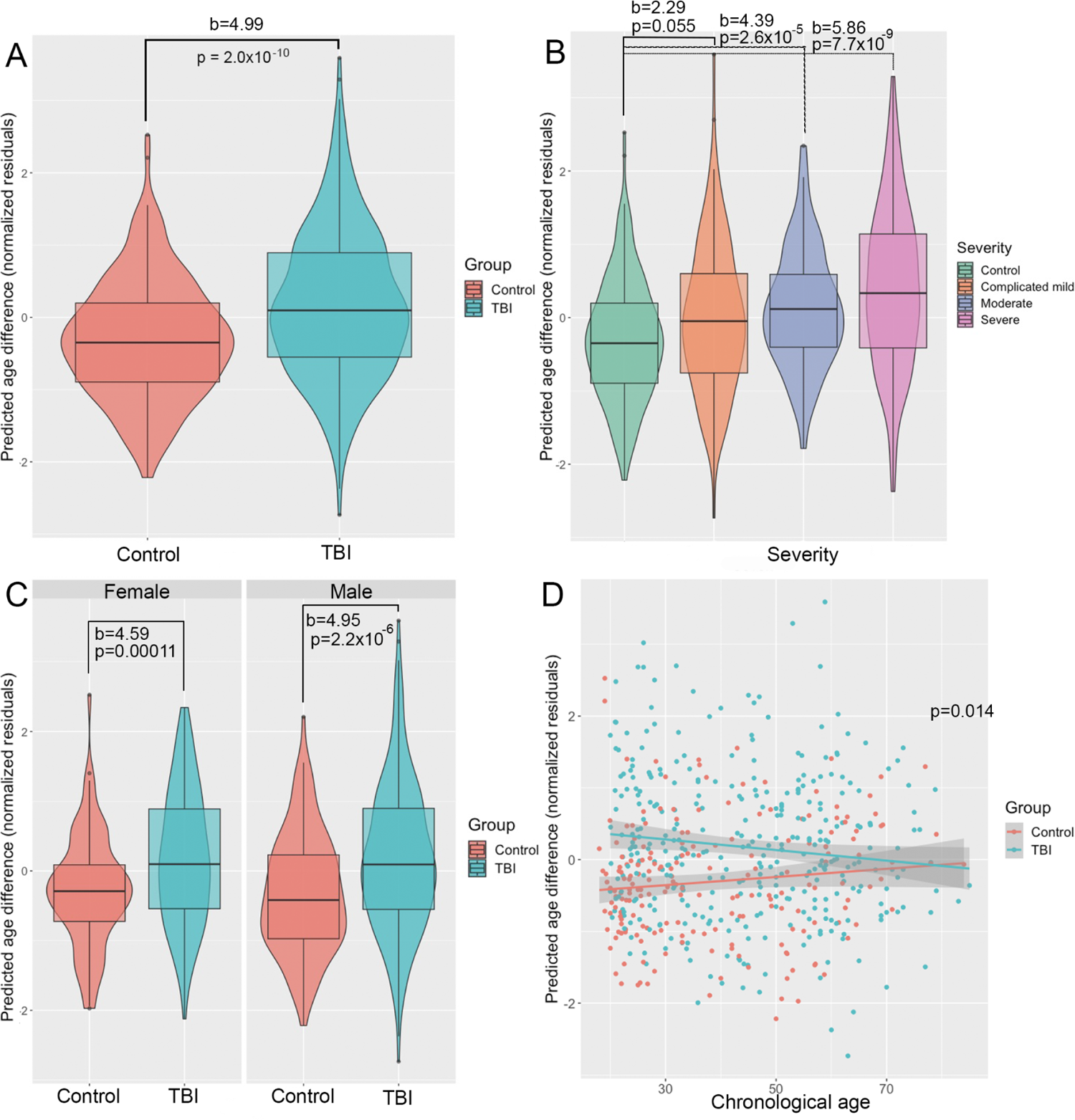
Group differences in PAD. (A), associations with severity (B), group × sex interaction (C), and group × age interaction (D). Box/violin plots are shown for group differences in panels A-C, with control group in red and cmsTBI group in blue (in A, C, and D). Statistics are displayed for each panel. Trendlines in panel D are linear estimates with 95% confidence intervals in gray.

#### Sensitivity analyses

The group difference in PAD remained after repeating the analysis while excluding participants with lesions visible on T1 scans (N=382, *b*=4.55, *p*=6.6×10^-8^) and excluding participants <1 year post-injury (N=476, *b*=5.02, *p*=3.7×10^-10^). Group differences were larger when individuals >60 years (age-at-scan) were excluded (N=458, *b*=5.95, *p*=8.5×10^-12^) and slightly smaller when patients injured as children were included (N=515, *b*=4.69, *p*=1.3×10^-9^).

### Within-TBI analyses

#### TSI

Across all cmsTBI participants more than a year post-injury, there was no linear association between TSI and PAD (N=276, *b*=0.14, *p*=.10), but there was a nonlinear association (*b*=0.02, *p*=.04). There was however, a positive linear association between TSI and PAD across cmsTBI participants more than five years post-injury (N=109, *b*=0.25, *p*=.02) and a non-significant trend in participants more than ten years post-injury (N=58, *b*=0.34, *p*=.057), but no nonlinear associations in either of these age brackets (**Figure 3**). Fitting natural splines with 3, 5, or 7 degrees of freedom (DF), we found that both the 3 and 5 DF spline models were significant (*p*s<.05) but that the 5 DF model yielded a slight improvement in model fit (based on Akaike Information Criterion). These curves exhibit an initial increase in brain age, followed by a decrease, followed by a slow and continuous increase (**Figure 3**).

**Figure 3.**
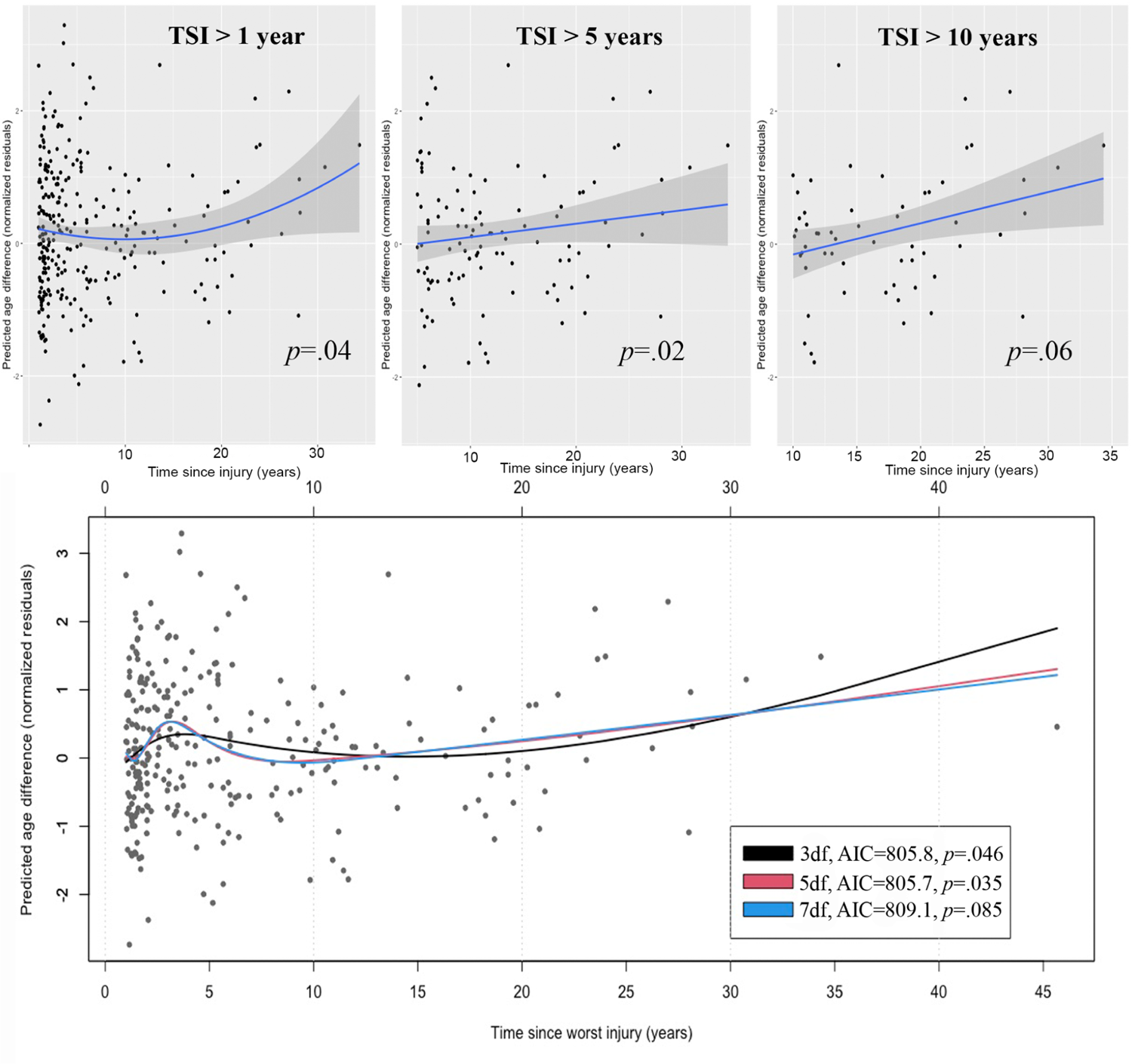
Association between PAD and TSI. The associations between time since injury in years and PAD (normalized residuals - accounting for age, sex, and random effects of cohort | site) are shown for three post-injury intervals (greater than 1 year, greater than 5 years, and greater than 10 years). Trendlines were plotted in R 4.2.2 with 95% confidence intervals in gray. There was a significant non-linear relationship with TSI across all cmsTBI participants more than 1 year post-injury and linear relationships with TSI beginning at 5 and 10 years post-injury. Given the significant nonlinear association, we also added a spline interpolation to better understand the shape. Using the *gam* function in the *splines* R library, we fit a natural spline with 3, 5, and 7 degrees of freedom (DF). The model with 5 DF provided the best fit. AIC=Akaike Information Criterion.

#### Age

The positive association between TSI and PAD did not hold for the individuals who were older at age-of-scan, in fact it was reversed (**Supplementary** Figure 4). This inversion, starting around age 65, suggests that survivor bias may have influenced brain age results in the oldest individuals. There are many factors that can influence longevity, most of which were not accessible with the data available, with the exception of education. This possibility of survivor bias in our data is supported by the greater portion of older individuals (>60 years at age-at-injury) who were highly educated (Bachelor’s degree and up, ISCED level 6+) (χ^2^ [6, *N*=518] = 14.2, *p*=.028; **Supplementary** Figure 7 and **Supplementary Table 4**). Education is a well-known proxy for cognitive reserve that may mitigate some aging effects.

#### Cognitive performance

There were no significant associations between PAD and any of the cognitive tests included across the cmsTBI group (DSF: N=175, *p*=.48; DSB: N=175, *p*=.29; TMT-A: N=173, *p*=.43; TMT-B: N=173, *p*=.86). These models were also non-significant when we covaried for education or tested associations across the full sample.

### Interactions

There was a significant group × sex interaction (*b*=1.63, *p*=.009, **Figure 2c**), indicating that group differences in PAD (cmsTBI vs Controls) were not the same between males and females. Group differences among females/women (*b*=4.59, N=179) were slightly smaller than among males/men (*b*=4.95, N=297). These remained significant after covarying for TSI or severity. There was also a significant group × age interaction (*b*=-0.11, *p*=.014, **Figure 2d**), indicating that the group differences were not consistent over adulthood. In fact, group differences were smaller among older participants. There were no significant interactions between chronological age × TSI, TSI × sex, or TSI × education in the cmsTBI group.

### APOE analyses

We hypothesized that individuals with at least one APOE ε4 allele would have greater PAD. The analysis included 56 individuals hetero- or homozygous for the ε4 allele and 110 individuals without an ε4 allele. The hypothesis was not supported; there was no significant difference in PAD based on APOE genotype in the whole sample or within the cmsTBI group (*p*=.64 and .48, respectively). There were also no significant interactions between APOE × group or APOE × age (*p*=.97 and .40, respectively).

## Discussion

Our goal was to examine the effects of remote cmsTBI on brain aging. In a mega-analysis of 540 individuals from 7 cohorts, we show evidence of accelerated brain aging after cmsTBI. Individuals with cmsTBI showed a PAD five years greater than the control group, consistent with other data in cmsTBI.^13,15^ These analyses extend prior work in three ways. First, the current sample included a large number of younger and older adults with a range of post-injury intervals. While prior work has focused on young to middle aged adults in the first few years after injury (e.g., mean 28 months post-injury^13^) or older adults in the late chronic timepoint (over 10 years post-injury),^15^ the median age for this study was 41.2 (range 18-85) and a mean TSI of 5.3±6.4 years (range 0.2-34 years) with 198 participants at least 50 years of age (age-at-scan, mean TSI 7.5±7.4 years). Second, there have been mixed findings with respect to the effects of injury severity and brain age, which are addressed below. Finally, in view of the established role of sex in TBI recovery, the current sample size allowed us to examine potential sex differences.

### Brain age and TSI

The positive association between TSI and PAD supports an accelerated aging effect, with the most dramatic effects observed after a decade post-injury (**Figure 3**). These data are consistent with prior work and extend those findings to a longer post-injury period. Spline interpolation revealed a pattern of initial injury response and an upward inflection in brain age (1-3 years in our data), followed by a slight decrease in brain age, perhaps reflecting injury accommodation and compensation, followed by a progressing increase in PAD around 7 years post-injury. While the year markers reported are biased by the TSI distribution in our sample (weighted towards TSI<10 years), we present the general shape and progression, which mirrors a biphasic response to injury observed in other measurements of TBI pathology including serum protein concentration.^31^

There remain a number of possible interacting variables including blood-brain barrier disruption and neuroinflammation^32^, or functional network changes that facilitate functional recovery in the acute stage but may promote proteinopathy over the long-term.^33^ Our data are consistent with the idea that brain volumetric changes post-injury cannot be accounted for by lesion resolution and transsynaptic or Wallerian events, and are likely attributable to more insidious physiological processes post-injury. While brain age may serve as a sensitive marker for brain health, future research efforts should clarify the clinical significance of this variable, establish the critical windows when cortical atrophy is occurring, and define the clinical and demographic modifiers of post-traumatic increase in PAD as well as its underlying mechanisms.

### Injury severity and PAD

In contrast to others,^2,17,18^ Cole and colleagues found little evidence of PAD in a single uncomplicated mild TBI event. On the other hand, a pronounced effect has been seen in cmsTBI.^13^ With the benefit of a larger cmsTBI sample, the current results extend these findings, showing a stepwise increase in PAD with injury severity (**Figure 3**). This finding is intuitive and consistent with extensive literature supporting a direct relationship between injury severity and atrophy based on T1w-MRI volumetrics.^28^

### Sex and PAD

There is growing evidence that sex plays a critical role in TBI recovery and outcome.^22^ The reasons for observed sex-based disparities in TBI outcome are a growing area of investigation, a welcome transition after decades of focus on male-only models for TBI.^34^ While some evidence points to hormonal differences as a factor contributing,^35^ mechanisms are still poorly understood. One aspect complicating sex comparisons is that the mechanism of injury may differ in ways that have implications for outcome.^36^ With regard to sex and brain morphometry, recent work in mild TBI revealed no differences.^37^ Our findings point to modestly greater PAD in men, although the effect was small (**Figure 2c**).

### Brain age and genetics

Contrary to our hypotheses, APOE ε4 did not have a greater PAD compared to non-carriers. Of note, the APOE literature is mixed, with some findings revealing a clear effect of APOE on brain volumetrics,^38^ while others show no relationship.^39^ In one recent examination of over 1100 individuals, examiners showed a differential effect of APOE on brain volume with the ε4 allele based on age, i.e. a paradoxical “protective” effect in individuals <60 years of age^40^ that reversed in older age groups. The null finding in the present data could *not* be explained by greater injury severity in the non-carriers or sex differences between subgroups. This null finding should be interpreted cautiously due to limited statistical power, but may also point to limitations in brain structure-based analysis in assessing functional changes; where brain organization, including functional connectivity, may better approximate neurological resilience.^41^ However, this null finding is consistent with a large genetic association study in over 4700 patients with TBI, which did not replicate the effect of APOE ε4 carrier status on outcome.^42^

### Brain age and behavioral outcome

We observed no significant relationship between the accelerating PAD in cmsTBI and cognitive functioning, i.e. psychomotor speed, executive functioning, and working memory. This finding is inconsistent with some prior work,^2,13^ but generally in line with a long history of incongruence between behavior and brain structure,^43^ given the numerous factors responsible for change in brain volume and for behavioral decline, including the role of neural and cognitive reserves. Therefore, while we did not find cognitive consequences for advancing PAD, this negative result may be simply due to the fact that neural networks deteriorate prior to the appearance of clinical changes. In support of this, a recent study showed PAD to be predictive of later progression to dementia in a typical aging sample.^44^

### Addressing challenges to external validity

The study of cmsTBI over the lifespan poses natural challenges to study design and subject enrollment. Subject recruitment in cmsTBI has multiple challenges, including sample bias with regard to socioeconomic status, sex and race.^45^ Survivor bias can also impact studies of illness, aging, and mortality, and may amplify counterintuitive effects.^46^

In our data, survivor bias may have contributed to the finding that PAD decreases with greater TSI in the oldest individuals (>65 years of age). Survivor bias is often difficult to track and demonstrate, but we used uncharacteristic differences in the sample as markers for potential biased sampling. Education is a predictor of longevity, and a proxy for cognitive reserve, which may serve to increase resilience post-cmsTBI.^47^ In our data, there were more individuals with advanced degrees (Bachelor’s and higher) beginning at age 60. One explanation for this upward inflection in sample education is age-related attrition in less educated individuals (e.g., due to mortality and dementia^48^) and greater health and resilience factors associated with advanced education, which is a modifier of MRI brain volume.^49^ *Aging with TBI* is distinct from *TBI during aging,* partly because TBIs sustained in older individuals are more likely due to falls (lower impact) than motor vehicles (higher impact) producing different brain pathology. Moreover, individuals who sustain a *TBI during aging (older age-at-injury)*, will typically have completed their educational goals and potential, whereas those who are *aging with TBI (younger age-at-injury)* may have their educational and career trajectories negatively altered by TBI.

Cross-sectional research addressing questions that are by nature developmental or evolve over a lifespan have significant limitations, and is the most important limitation of our study. To ideally address the goals of the current study, large datasets with longitudinal data spanning decades are needed. We also recognize that a majority of the sample was from participants of white-European descent, and there is a great need to determine the role of health care access, race, and socioeconomic status on outcomes.^45^ This is particularly true given that non-whites are less likely to be represented in the research literature,^45^ have poorer clinical outcomes post-TBI,^45^ and potentially carry a higher risk for Alzheimer’s disease and related dementias.^50^ Finally, while the methodology chosen in this study to determine brain age is well validated,^13,14^ there are multiple additional brain age algorithms that could also be used.^8^

## Conclusions

The current mega-analysis reveals that a single cmsTBI is associated with a PAD of nearly 5 years along with *accelerated aging* years after injury. We show a non-linear time course for the initial effects of injury, followed by relative stability, and then faster progression of brain age in the late chronic phase. Injury severity showed a stepwise relationship with advancing brain age. There was also a mild influence of sex, with men showing relatively larger PAD. Longitudinal designs are needed to assess disease progression or mitigation. Brain age holds promise as a useful biomarker to track changes over time due to its dynamic nature and amenability to modification with preventive measures (e.g., lifestyle adjustments) and, possibly, treatment.

## Acknowledgements

NIH/NINDS R61NS120249 to Dr. Dennis. Dr. Menon is an Emeritus Senior Investigator of the National Institute for Health Research, UK, and is supported by the CENTER-TBI grant from the European Union and the TBI-REPORTER grant from UKRI, NIHR, AR UK and UK Ministry of Defense. Israel Innovation Authority, “Magneton” Grant to Dr Livny. Maryland Technology Development Corporation1 R01 EY028039 Congressionally Directed Medical Research, JH Discovery Award to Dr. Koliatsos. National Institute for Health and Care Research (NIHR) Advanced Fellowship to Dr. Newcombe. NHMRC Early Career Fellowship to Dr. Spitz. NIH grants R01NS100973, R01AG079957 and RF1AG082201, DoD contract W81XWH-18-1-0413, an anonymous donor family, and the Hanson-Thorell Research Scholarship Fund at the University of Southern California to Dr. Irimia. NIH/NIA R01AG061028, NIH/NINDS RF1NS115268, NIH/NINDS RF1NS128961 and NIH/NINDS/NICHD1U01NS086625-01 to Dr. Dams-O’Connor. NIH/NINDS R01-NS121107 to Dr. Dobryakova. PA Health Research Grant SAP #4100077082 to Dr. Hillary. The European Research Council (ERC) under the European Union’s Horizon 2020 research and innovation programme (ERC Starting Grant, Grant agreement No. 802998) and the Research Council of Norway (249795) to Dr. Westlye. To Dr. Caeyenberghs: The Victorian Near-miss Award Pilot is administered by veski for the Victorian Health and Medical Research Workforce Project on behalf of the Victorian Government and the Association of Australian Medical Research Institutes. Funding for the Pilot has been provided by the Victorian Department of Jobs, Precincts and Regions. The Liaison Committee between the Central Norway Regional Health Authority (RHA) and the Norwegian University of Science and Technology (NTNU) (2020/39645) to Dr. Olsen. Tiny Blue Dot Foundation to Dr. Monti. Transport Accident Commission to Dr. Ponsford. NIH U54EB020403, R01MH116147, R56AG058854, P41EB015922, R01MH111671 to Dr. Thompson. VA R&D SPIRE AWARD.

## Author contributions

ELD, MMA, HA, EDB, KC, JHC, KDOC, EMD, ED, HMG, JHG, AKH, TH, AI, VEK, HML, AL, DKM, TLM, AZM, SM, MMM, VFJN, MRN, JP, HS, GS, LTW, RZ, PMT, EAW, AO, and FGH contributed to the conception and design of the study. ELD, SV, MMA, KC, JHC, KDOC, ED, HMG, TH, AR, GS, UV, LTW, AO, and FGH contributed to the acquisition and analysis of data. ELD and FGH contributed to drafting the text of preparing the figures.

## References

1. Graham NS, Sharp DJ. Understanding neurodegeneration after traumatic brain injury: from mechanisms to clinical trials in dementia. J. Neurol. Neurosurg. Psychiatry 2019;90(11):1221–1233.

2. Amgalan A, Maher AS, Ghosh S, et al. Brain age estimation reveals older adults’ accelerated senescence after traumatic brain injury. Geroscience 2022;44(5):2509–2525.

3. Graham NSN, Cole JH, Bourke NJ, et al. Distinct patterns of neurodegeneration after TBI and in Alzheimer’s disease. Alzheimers. Dement. 2023;19(7):3065–3077.

4. Lu Y, Jarrahi A, Moore N, et al. Inflammaging, cellular senescence, and cognitive aging after traumatic brain injury. Neurobiol. Dis. 2023;180:106090.

5. Huang X, You W, Zhu Y, et al. Microglia: A Potential Drug Target for Traumatic Axonal Injury. Neural Plast. 2021;2021:5554824.

6. Ayubcha C, Revheim M-E, Newberg A, et al. A critical review of radiotracers in the positron emission tomography imaging of traumatic brain injury: FDG, tau, and amyloid imaging in mild traumatic brain injury and chronic traumatic encephalopathy. Eur. J. Nucl. Med. Mol. Imaging 2021;48(2):623–641.

7. Moretti L, Cristofori I, Weaver SM, et al. Cognitive decline in older adults with a history of traumatic brain injury. Lancet Neurol. 2012;11(12):1103–1112.

8. Yin C, Imms P, Cheng M, et al. Anatomically interpretable deep learning of brain age captures domain-specific cognitive impairment. Proc. Natl. Acad. Sci. U. S. A. 2023;120(2):e2214634120.

9. Franke K, Gaser C. Ten Years of BrainAGE as a Neuroimaging Biomarker of Brain Aging: What Insights Have We Gained? Front. Neurol. 2019;10:789.

10. Han LKM, Dinga R, Leenings R, et al. A large-scale ENIGMA multisite replication study of brain age in depression. Neuroimage: Reports 2022;2(4):100149.

11. Clausen AN, Fercho KA, Monsour M, et al. Assessment of brain age in posttraumatic stress disorder: Findings from the ENIGMA PTSD and brain age working groups. Brain Behav. 2022;12(1):e2413.

12. Liew S-L, Schweighofer N, Cole JH, et al. Association of Brain Age, Lesion Volume, and Functional Outcome in Patients With Stroke [Internet]. Neurology 2023;Available from: 10.1212/WNL.0000000000207219

13. Cole JH, Leech R, Sharp DJ, Alzheimer’s Disease Neuroimaging Initiative. Prediction of brain age suggests accelerated atrophy after traumatic brain injury. Ann. Neurol. 2015;77(4):571–581.

14. Cole JH, Jolly A, de Simoni S, et al. Spatial patterns of progressive brain volume loss after moderate-severe traumatic brain injury. Brain 2018;141(3):822–836.

15. Spitz G, Hicks AJ, Roberts C, et al. Brain age in chronic traumatic brain injury. Neuroimage Clin 2022;35:103039.

16. Newcombe VFJ, Ashton NJ, Posti JP, et al. Post-acute blood biomarkers and disease progression in traumatic brain injury. Brain 2022;145(6):2064–2076.

17. Dennis EL, Taylor BA, Newsome MR, et al. Advanced brain age in deployment-related traumatic brain injury: A LIMBIC-CENC neuroimaging study. Brain Inj. 2022;1–11.

18. Gan S, Shi W, Wang S, et al. Accelerated Brain Aging in Mild Traumatic Brain Injury: Longitudinal Pattern Recognition with White Matter Integrity. J. Neurotrauma 2021;38(18):2549–2559.

19. Poudel GR, Dominguez D JF, Verhelst H, et al. Network diffusion modeling predicts neurodegeneration in traumatic brain injury. Ann Clin Transl Neurol 2020;7(3):270–279.

20. Thompson PM, Jahanshad N, Ching CRK, et al. ENIGMA and global neuroscience: A decade of large-scale studies of the brain in health and disease across more than 40 countries. Transl. Psychiatry 2020;10(1):100.

21. Olsen A, Babikian T, Bigler E, et al. Toward a Global and Open Science for Imaging Brain Trauma: the ENIGMA Adult msTBI Working Group. Brain Imaging Behav. 2019;Under Review

22. Gupte R, Brooks W, Vukas R, et al. Sex Differences in Traumatic Brain Injury: What We Know and What We Should Know. J. Neurotrauma 2019;36(22):3063–3091.

23. Maiti TK, Konar S, Bir S, et al. Role of apolipoprotein E polymorphism as a prognostic marker in traumatic brain injury and neurodegenerative disease: a critical review. Neurosurg. Focus 2015;39(5):E3.

24. for Statistics UI. International standard classification of education: ISCED 2011 [Internet]. Comp. Soc. Res. 2012;30Available from: https://www.voced.edu.au/content/ngv:54992

25. Hacker D, Jones CA, Yasin E, et al. Cognitive Outcome After Complicated Mild Traumatic Brain Injury: A Literature Review and Meta-Analysis [Internet]. J. Neurotrauma 2023;Available from: 10.1089/neu.2023.0020

26. Karatzoglou A, Smola A, Hornik K, Zeileis A. Kernlab - an S4 package for kernel methods in R. J. Stat. Softw. 2004;11(9):1–20.[cited 2021 Mar 18]

27. de Lange A-MG, Anatürk M, Rokicki J, et al. Mind the gap: Performance metric evaluation in brain-age prediction. Hum. Brain Mapp. 2022;43(10):3113–3129.

28. Bigler ED. Distinguished Neuropsychologist Award Lecture 1999. The lesion(s) in traumatic brain injury: implications for clinical neuropsychology. Arch. Clin. Neuropsychol. 2001;16(2):95–131.

29. McHutchison CA, Backhouse EV, Cvoro V, et al. Education, Socioeconomic Status, and Intelligence in Childhood and Stroke Risk in Later Life: A Meta-analysis. Epidemiology 2017;28(4):608–618.

30. Deary IJ, Taylor MD, Hart CL, et al. Intergenerational social mobility and mid-life status attainment: Influences of childhood intelligence, childhood social factors, and education. Intelligence 2005;33(5):455–472.

31. Shahim P, Politis A, van der Merwe A, et al. Time course and diagnostic utility of NfL, tau, GFAP, and UCH-L1 in subacute and chronic TBI. Neurology 2020;95(6):e623–e636.

32. Visser K, Koggel M, Blaauw J, et al. Blood-based biomarkers of inflammation in mild traumatic brain injury: A systematic review. Neurosci. Biobehav. Rev. 2022;132:154–168.

33. Hillary FG, Grafman JH. Injured Brains and Adaptive Networks: The Benefits and Costs of Hyperconnectivity. Trends Cogn. Sci. 2017;21(5):385–401.

34. Shansky RM, Murphy AZ. Considering sex as a biological variable will require a global shift in science culture. Nat. Neurosci. 2021;24(4):457–464.

35. Franke K, Hagemann G, Schleussner E, Gaser C. Changes of individual BrainAGE during the course of the menstrual cycle. Neuroimage 2015;115:1–6.

36. Mikolić A, van Klaveren D, Groeniger JO, et al. Differences between Men and Women in Treatment and Outcome after Traumatic Brain Injury. J. Neurotrauma 2021;38(2):235–251.

37. Shida AF, Massett RJ, Imms P, et al. Significant acceleration of regional brain aging and atrophy after mild traumatic brain injury [Internet]. J. Gerontol. A Biol. Sci. Med. Sci. 2023;Available from: 10.1093/gerona/glad079

38. Rogojin A, Gorbet DJ, Hawkins KM, Sergio LE. Differences in structural MRI and diffusion tensor imaging underlie visuomotor performance declines in older adults with an increased risk for Alzheimer’s disease. Front. Aging Neurosci. 2022;14:1054516.

39. Vervoordt SM, Arnett P, Engeland C, et al. Depression associated with APOE status and hippocampal volume but not cognitive decline in older adults aging with traumatic brain injury. Neuropsychology 2021;35(8):863–875.

40. Tang M, Su N, Zhang D, et al. The Differential Effects of Apoeɛ4 on Cerebral Volumetric Structures in Different Lifespan in Community-Dwelling Populations [Internet]. J. Alzheimers. Dis. 2023;Available from: 10.3233/JAD-220834

41. Stern Y, Varangis E, Habeck C. A framework for identification of a resting-bold connectome associated with cognitive reserve. Neuroimage 2021;232:117875.

42. Kals M, Kunzmann K, Parodi L, et al. A genome-wide association study of outcome from traumatic brain injury. EBioMedicine 2022;77:103933.

43. Marek S, Tervo-Clemmens B, Calabro FJ, et al. Reproducible brain-wide association studies require thousands of individuals. Nature 2022;603(7902):654–660.

44. Biondo F, Jewell A, Pritchard M, et al. Brain-age is associated with progression to dementia in memory clinic patients. Neuroimage Clin 2022;36:103175.

45. Brenner EK, Grossner EC, Johnson BN, et al. Race and ethnicity considerations in traumatic brain injury research: Incidence, reporting, and outcome. Brain Inj. 2020;34(6):799–808.

46. Mitchell E, Chohan H, Bestwick JP, Noyce AJ. Alcohol and Parkinson’s Disease: A Systematic Review and Meta-Analysis. J. Parkinsons. Dis. 2022;12(8):2369–2381.

47. Stern Y, Barnes CA, Grady C, et al. Brain reserve, cognitive reserve, compensation, and maintenance: operationalization, validity, and mechanisms of cognitive resilience. Neurobiol. Aging 2019;83:124–129.

48. Fuller GW, Ransom J, Mandrekar J, Brown AW. Long-Term Survival Following Traumatic Brain Injury: A Population-Based Parametric Survival Analysis. Neuroepidemiology 2016;47(1):1–10.

49. Steffener J. Education and age-related differences in cortical thickness and volume across the lifespan. Neurobiol. Aging 2021;102:102–110.

50. Barnes LL. Alzheimer disease in African American individuals: increased incidence or not enough data? Nat. Rev. Neurol. 2022;18(1):56–62.

