## Supplement for "Accelerated Aging after Traumatic Brain Injury: an ENIGMA Multi-Cohort Mega-Analysis"

**Supplementary Materials**

**Supplementary Table 1. Inclusion/exclusion criteria by site.**

**Supplementary Table 2. Scan parameters by site.**

**Supplementary Table 3. Exclusion reasons by site.**

**Supplementary Figure 1. Chronological age versus brain age by site.**

**Supplementary Figure 2. Statistical model flowchart.**

**Supplementary Figure 3. Injury severity binned by age at injury.** 1=complicated mild TBI, 2=moderate TBI, 3=severe TBI.

**Supplementary Figure 4. Effects of time since injury on PAD separated by age category.** Older = over 65 years old at time of injury, younger = under 65 years old at time of injury.

**Supplementary Figure 5. PAD separated by age brackets - age at scan (upper) and age at injury (lower).**

**Supplementary Figure 6. Residual associations with chronological age within TSI bins.**

**Supplementary Figure 7. Educational levels broken down by age at injury bracket.**

**Supplementary Table 4. Demographics broken down by age at injury.** For each decade, the total N, % female, education level (ISCED 2011[1]), and severity are shown. NA=not available, cmTBI=complicated mild TBI, modTBI=moderate TBI, sevTBI=severe TBI.

**Supplementary Table 1. Inclusion/exclusion criteria by site.**

| Cohort | Recruitment | Inclusion criteria | Exclusion criteria | How severity was determined |
|---|---|---|---|---|
| Kessler | Participants were recruited from Kessler Foundation database and the local community | 1) non-penetrating msTBI (moderate-severe) (intake or post-resuscitation Glasgow Coma Scale (GCS) score between 3 and 12); 2) 18-65 years of age; 3) right-handed; 4) normal visual acuity or vision corrected with contact lenses/eyeglasses; and 5) English skills sufficient to understand instructions and be familiar with common words (the neuropsychological tests used in this study presume competence in English). | 1) history of neurological illness, such as prior msTBI, brain tumor or severe seizures; 2) motor deficits that prevent the subject from being examined in an MR scanner (e.g., spasms, movement disorder); 3) history of psychosis, ADHD, Tourette's Disorder, learning disability, mental retardation, autism, or substance abuse. These conditions are associated with cognitive impairments that might overlap with those caused by TBI. MRI contraindication was also an exclusion criterion. | GCS and/or acute CT |
| LETBI | Participants were recruited from through Mt. Sinai inpatient rehabilitation records, local community and government agencies and advocacy groups, and social media. | 1) for TBI group: medically documented TBI or history of TBI reported on BISQ resulting in a period of altered mental status of loss of consciousness; 2) at least 18 years of age; 3) at least 1 year post injury; 4) able to give consent or have proxy give consent. | 1) participants who do not sign an autopsy intent form. | Medically documented TBI or history of TBI reported on BISQ resulting in a period of altered mental status of loss of consciousness |
| Monash | Individuals were recruited from admissions to a TBI rehabilitation center in the context of a no-fault accident compensation system. | Medically-confirmed non-penetrating TBI. English skills sufficient to understand instructions and be familiar with common words (the neuropsychological tests used in this study presume competence in English). At least 16 years of age. | Previous history of TBI or other neurological disorder. | GCS and/or, PTA, and/or acute CT |
| NTNU | Patients were recruited from a prospective database of admitted patients to the Department of Neurosurgery, St. Olav's Hospital, Trondheim University Hospital | for TBI group: non-penetrating msTBI (based on Head Injury Severity Scale) | 1) previous moderate or severe head injury; 2) diagnosed neurologic or psychiatric condition. Participants with contraindications to undergoing an MRI scan were excluded. | HISS |

| | | | | |
|---|---|---|---|---|
| Oslo | Patients were included in a prospective study comprising patients with acute mild traumatic brain injury admitted to Oslo University Hospital. | Mild traumatic brain injury here defined using the criteria from the American Congress of rehabilitation medicine, included patients aged 16-65 years with recent (<24 hours) history of trauma to the head (hospitalization with diagnosis S06.0-S06.9), resulting in loss of consciousness < 30 minutes, PTA< 24 hours and GCS between 13-15. The GCS was registered within the first 24 hours following injury, and the lowest GCS within the first 24 hours is reported. | Severe mental illness (e.g., major depressive disorder, schizophrenia or bipolar disorder diagnosed by a psychiatrist or clinical psychologist), progressive neurologic disease, previous ICD-10 diagnosis of substance dependence, contraindications for MRI (including pregnancy and claustrophobia), or lack of Norwegian language skills. | GCS and/or acute CT |
| PSU | Participants were recruited through Hershey Medical Center Departments of Neurology and PM&R, and from research registries at Moss Rehabilitation Research Institute in Philadelphia. | Moderate/severe TBI, defined as post-resuscitation GCS 3-12 or positive CT findings if GCS>12, documented loss of consciousness >30min, post-traumatic amnesia >24 hours. | PA-DOH: 410007708: at least 50 years of age and at least 5 years post-injury. Given that chronic traumatic encephalopathy has been observed in individuals as young as 45 years of age, in order to guarantee sensitivity to brain changes and possible degeneration we included individuals at least 50 years of age with no upper age restriction. For NIH Tr000127 and NJCBI 0120090178, age ranges included 18-75 years. For all studies, Individuals with significant neuropsychiatric disturbance (i.e., schizophrenia, bipolar disorder), or substance abuse requiring inpatient rehabilitation, or neurological disorder other than TBI were excluded. | GCS, PTA, time to follow commands, and/or acute CT |
| VA Palo Alto 1 | Participants (ages: 50-75) recruited through VA Palo Alto and community | History of mild-moderate TBI greater than 6 months from injury with residual memory.cognitive problems which interfere with daily functioning. | 1) Diagnosed with Dementia; 2) Pregnant or lactating female; 3) Unable to be safely withdrawn, at least two-weeks prior to beginning treatment, from medications that substantially increase the risk of seizures; 4) Have a cardiac pacemaker or a cochlear implant; 5) Have an implanted device (deep brain stimulation) or metal in the brain; 6) Have a mass lesion, cerebral infarct or other active CNS disease, including a seizure disorder; 7) Known current psychosis as determined by DSM-IV coding in chart (Axis I, psychotic disorder, schizophrenia) or a history of a non-mood psychotic disorder; 8) Diagnosis of Bipolar Affective Disorder I (as determined by chart review and intake interview), since this in conjunction with TBI increases seizure risk; 9) Current amnesic disorders, dementia, MOCA <16, or delirium.; 10) Current substance abuse (not including caffeine or nicotine) as determined by positive toxicology screen, or by history via AUDIT, within 3 months prior to screening; 11) Prior history of seizures; 12) Severe TBI or open head injury; 13) TBI within last 6 months; 14) Participation in another concurrent clinical trial; 15) Patients with prior exposure to rTMS (NOTE: TMS is allowed) or | OSU |

| | | | | |
|---|---|---|---|---|
| | | | ECT; 16) Active current suicidal intent or plan. Patients at risk for suicide will be required to establish a written safety plan involving their primary psychiatrist. All patients at risk for suicide will be excluded from the study (as per FDA recommendation). | |
| VA Palo Alto 2 | Participants over the age of 18 recruited from Veteran population seen in the War Related Illness and Injury Study Center | **Active Group:** Patients seen in the WRIISC clinical (local and national referrals), other VA and non-VA referring clinics. At least two or more of the following diagnosis: PTSD, depression, cognitive problems (including memory), skin rash, chronic pain, chronic GI problems.<br>**Control Group:** (see phone screen script and the questionnaire done at assessment) not a WRIISC patient referral, no diagnosis of PTSD, depression or cognitive and/or mental disorder no history of or current substance abuse/dependence | For 3T fMRI protocol:<br>Presence of; pacemakers, aneurysm clips, artificial heart valves, ear implants, metal fragments or foreign objects in the eyes, skin or body.<br>claustrophobia.<br>unable to understand (read/write/speak) English sufficiently to participate in informed consent hearing impairment which requires device/hearing aid<br>For the Control Group only - Axis I disorder<br>For EEG protocol:<br>ADHD, ADD, OCD, epilepsy, general seizure/seizure disorders, any movement disorder (e.g., Parkinson's, Tourette's); language disorders | OSU |

**Supplementary Table 2. Scan parameters by site.**

| Cohort | Scanner | Field strength | Voxel size |
|---|---|---|---|
| Kessler | Siemens Skyra | 3T | 1x1x1 |
| LETBI | Siemens Skyra | 3T | 1x1x1 |
| Monash | Siemens Verio | 3T | 1x1x1 |
| NTNU | Siemens Trio | 3T | 1x1x1 |
| Oslo | GE Signa HDxt | 3T | 1.2x1x1 |
| PSU | Siemens Prisma Fit | 3T | 1x1x1 |
| VA Palo Alto 1 | GE Discovery MR750 | 3T | 0.9x0.9x0.9 |
| VA Palo Alto 2 | GE Discovery MR750 | 3T | 0.6x1x1 |

**Supplementary Table 3. Exclusion reasons by site.**

| Cohort | Total excluded | QC | Outlier | Pediatric Injury | Uncomplicated mTBI | Missing clinical info |
|---|---|---|---|---|---|---|
| Kessler | 73 | 66 | 3 | 1 | 2 | 1 |
| LETBI | 25 | 6 | 0 | 16 | 3 | 0 |
| Monash | 21 | 9 | 1 | 3 | 8 | 0 |
| NTNU | 27 | 19 | 0 | 8 | 0 | 0 |
| Oslo | 5 | 0 | 0 | 5 | 0 | 0 |
| PSU | 43 | 25 | 6 | 3 | 6 | 3 |
| VA Palo Alto | 17 | 13 | 1 | 3 | 0 | 0 |
| Total | 211 | 138 | 11 | 39 | 19 | 4 |

**Supplementary Figure 1. Chronological age versus brain age by site.**

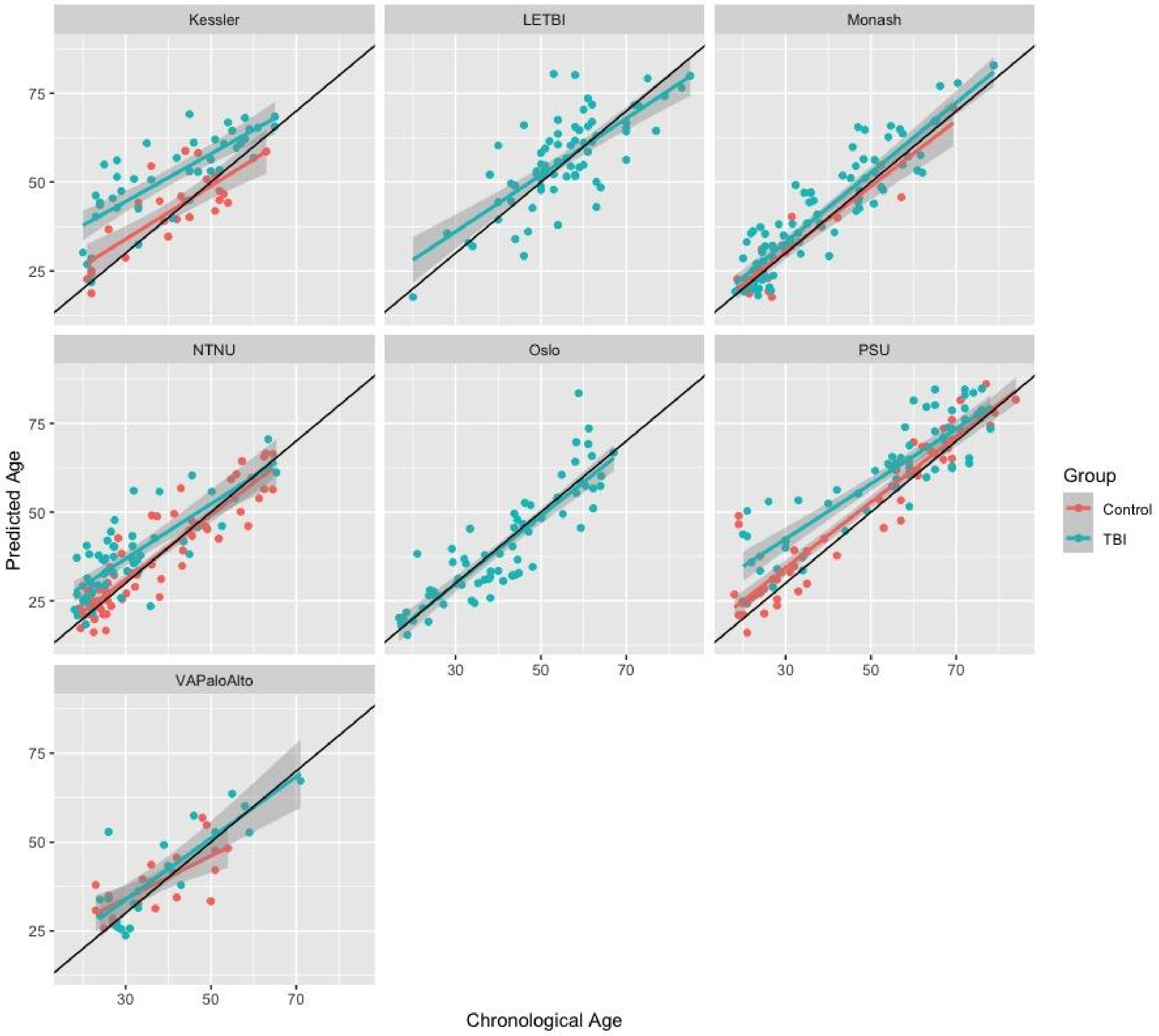

**Supplementary Figure 2. Statistical model flowchart.**

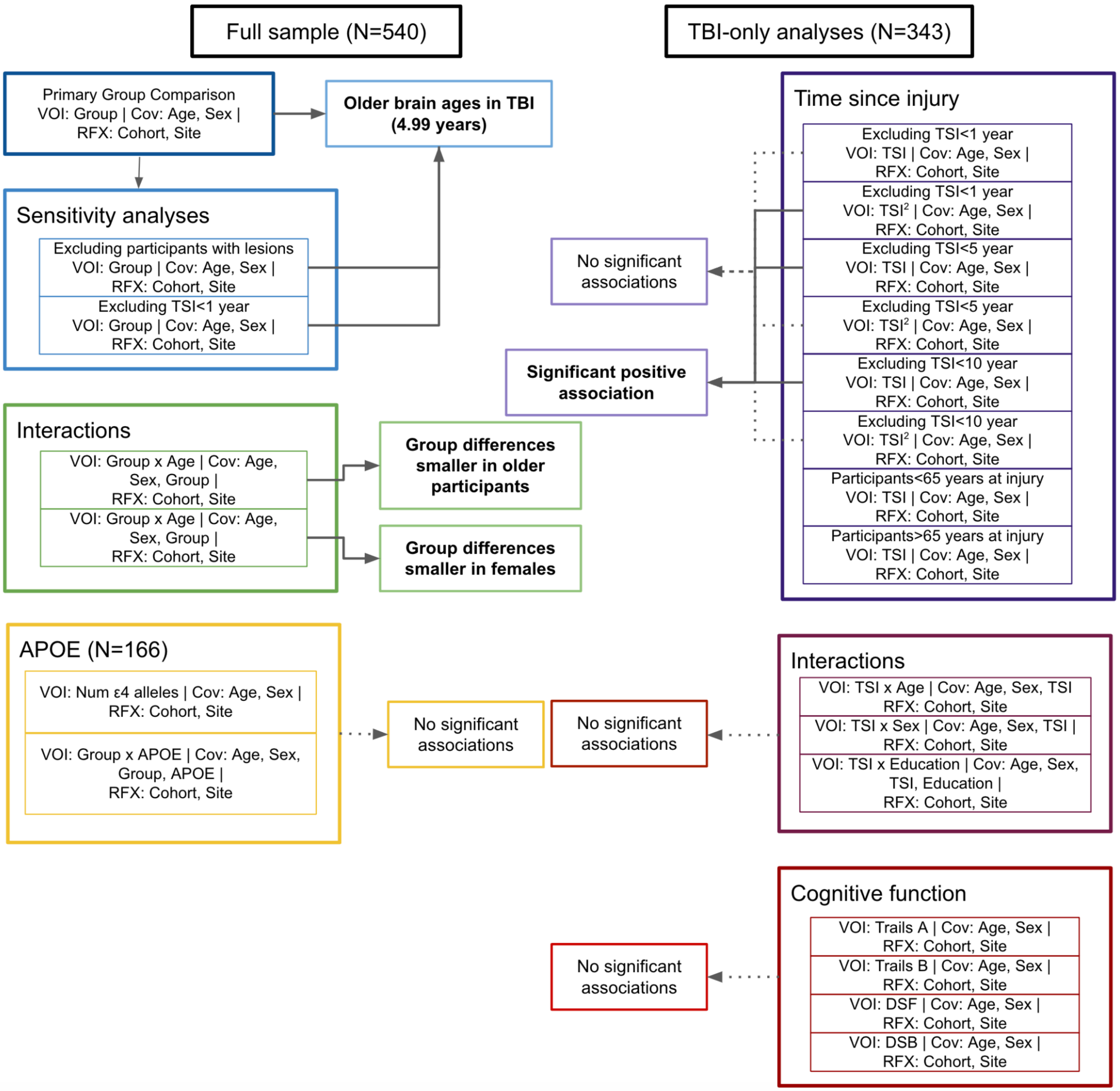

**Supplementary Figure 3. Injury severity binned by age at injury.** 1=complicated mild TBI, 2=moderate TBI, 3=severe TBI.

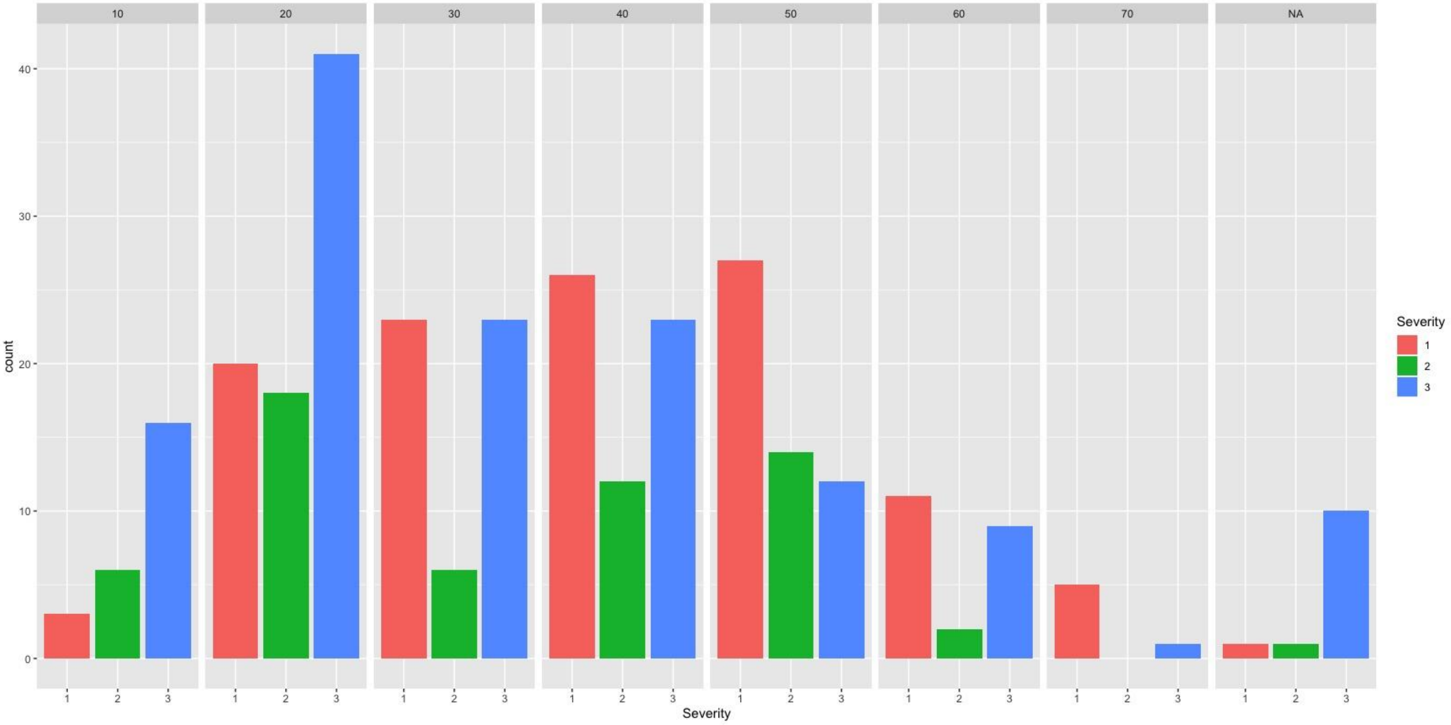

**Supplementary Figure 4. Effects of time since injury on PAD separated by age category.** Older = over 65 years old at time of injury, younger = under 65 years old at time of injury.

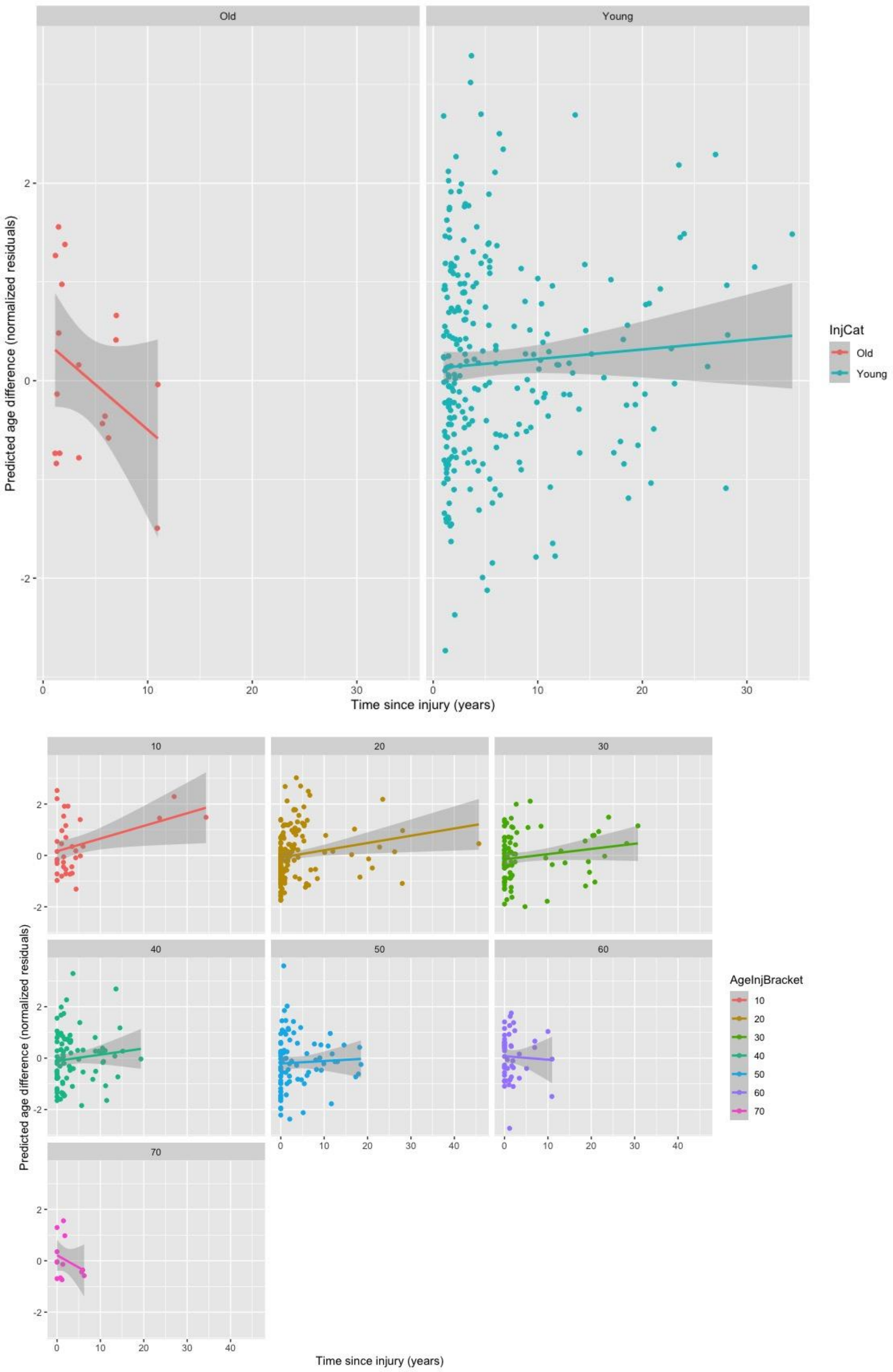

**Supplementary Figure 5. PAD separated by age brackets - age at scan (upper) and age at injury (lower).**

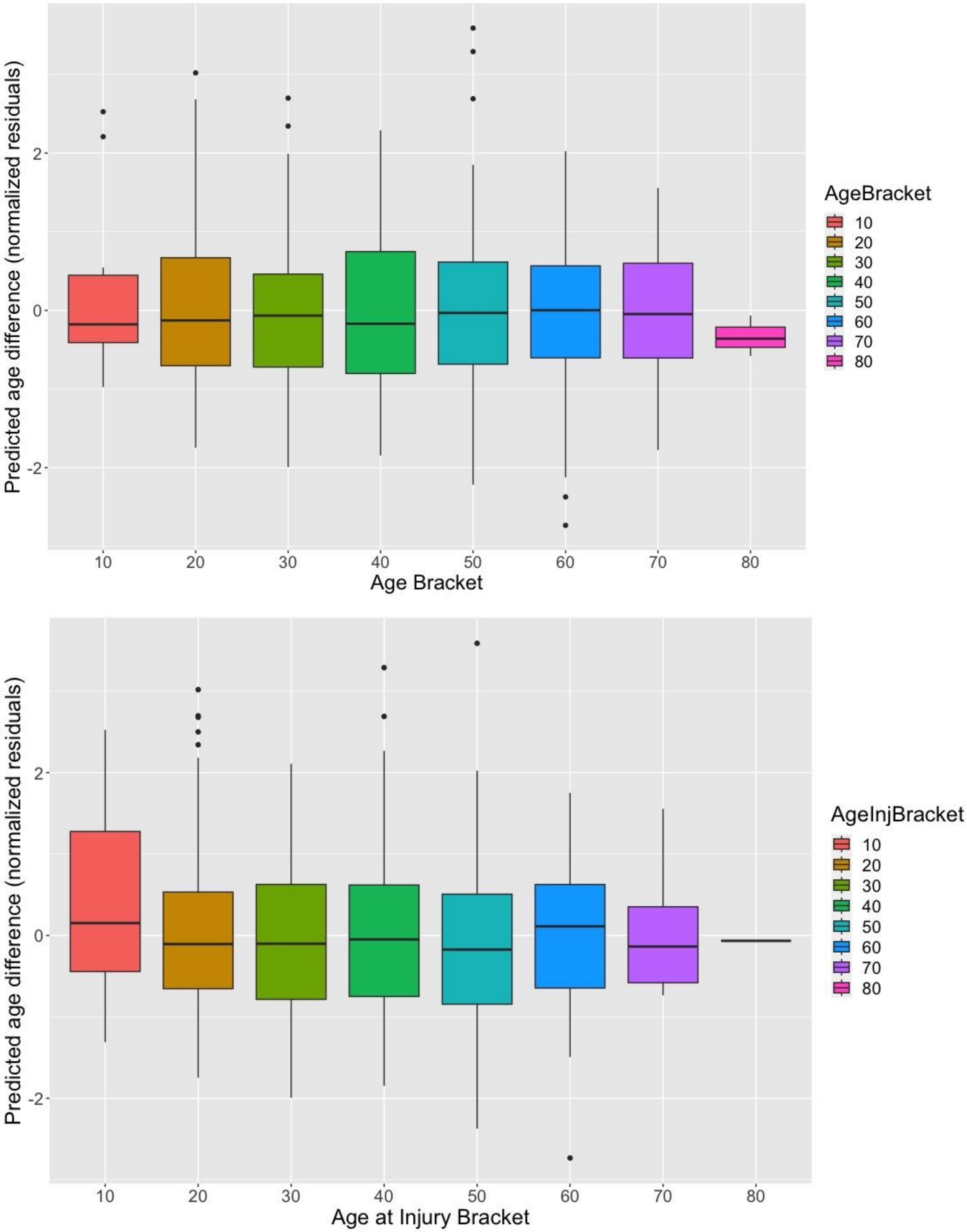

**Supplementary Figure 6. Residual associations with chronological age within TSI bins.**

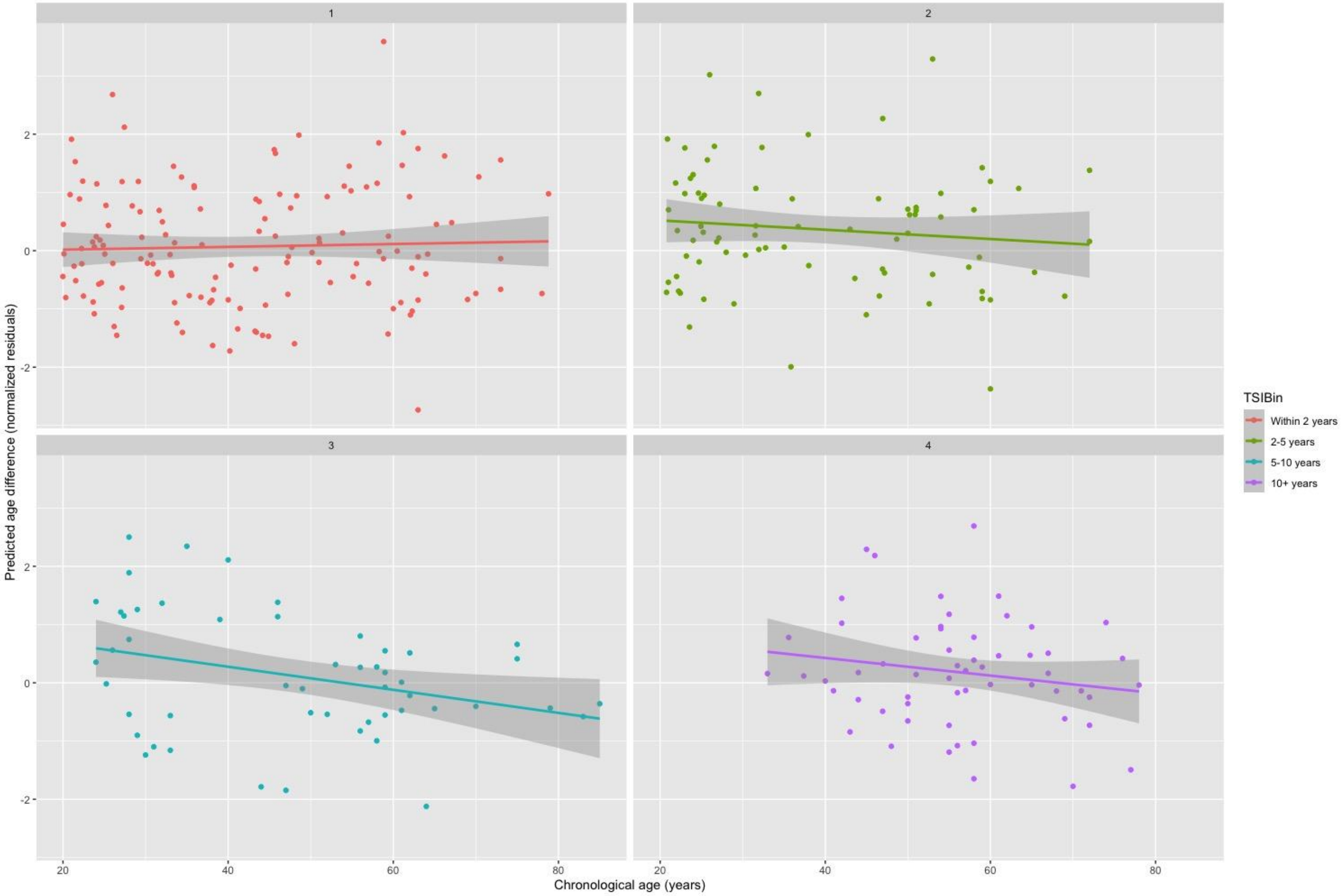

**Supplementary Figure 7. Educational levels broken down by age at injury bracket.** Proportion of sample broken down by ISCED 2011[1] categories are shown.

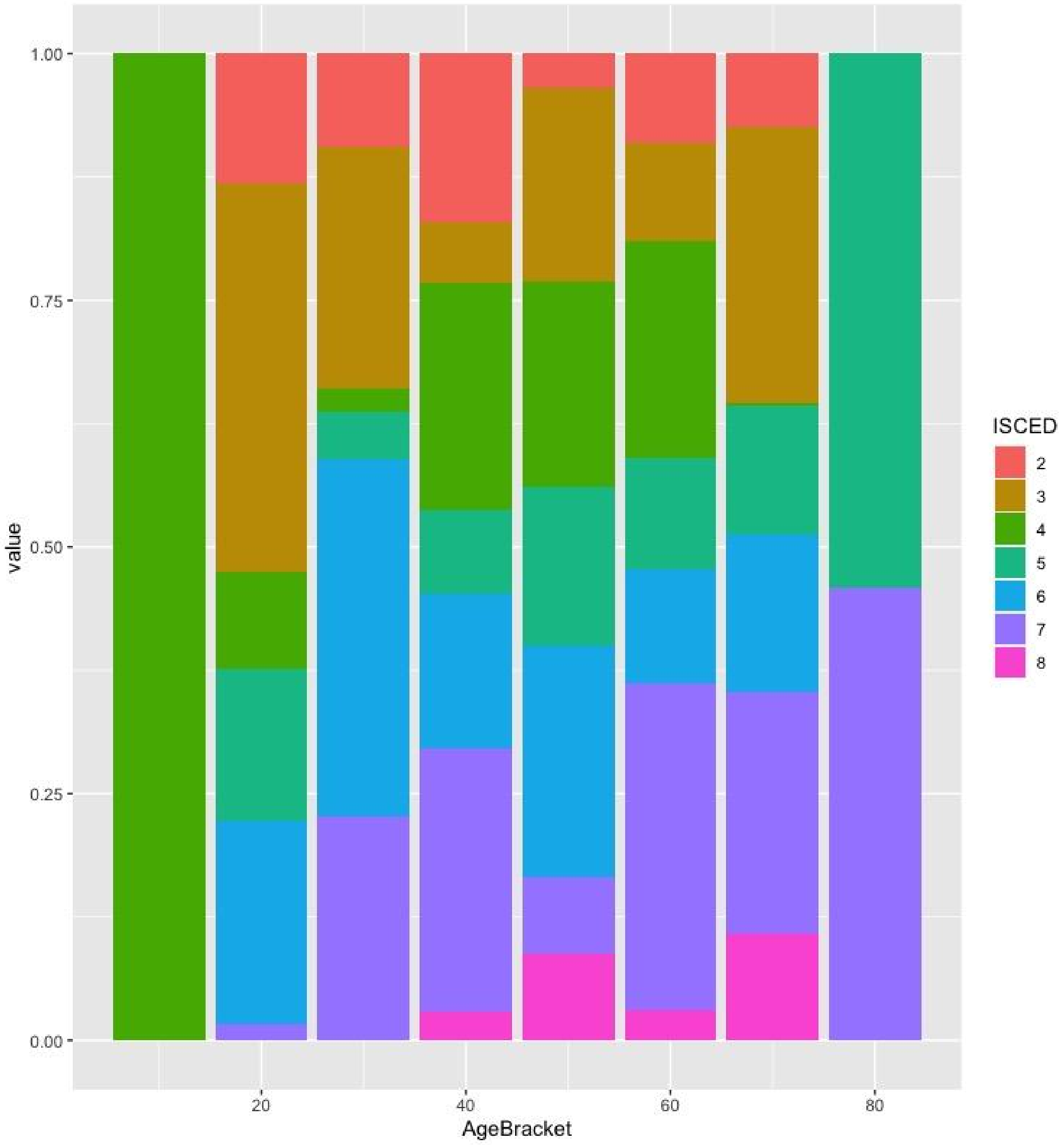

**Supplementary Table 4. Demographics broken down by age at injury.** For each decade, the total N, % female, education level (ISCED 2011[1]), and severity are shown. NA=not available, cmTBI=complicated mild TBI, modTBI=moderate TBI, sevTBI=severe TBI.

| | | Age at injury | | | | | | | |
|---|---|---|---|---|---|---|---|---|---|
| | | **Under 20** | **20-30** | **30-40** | **40-50** | **50-60** | **60-70** | **70-80** | **NA** |
| **N** | **Total** | 26 | 90 | 54 | 67 | 57 | 28 | 8 | 13 |
| | **% Female** | 23% | 27% | 31% | 33% | 32% | 50% | 25% | 23% |
| **ISCED (%)** | **2** | 15% | 10% | 9% | 13% | 5% | 18% | 12% | 8% |
| | **3** | 38% | 38% | 20% | 15% | 16% | 14% | 25% | 0% |
| | **4** | 19% | 7% | 6% | 13% | 21% | 18% | 0% | 8% |
| | **5** | 12% | 11% | 9% | 12% | 16% | 4% | 25% | 8% |
| | **6** | 15% | 23% | 37% | 21% | 18% | 14% | 12% | 15% |
| | **7** | 0% | 10% | 17% | 19% | 14% | 21% | 25% | 0% |
| | **8** | 0% | 0% | 0% | 6% | 9% | 7% | 0% | 0% |
| | **NA** | 0% | 1% | 2% | 0% | 2% | 4% | 0% | 0% |
| **Severity (%)** | **cmTBI** | 11% | 22% | 43% | 39% | 47% | 39% | 62% | 8% |
| | **modTBI** | 23% | 20% | 11% | 18% | 49% | 7% | 0% | 8% |
| | **sevTBI** | 61% | 46% | 43% | 34% | 21% | 32% | 12% | 77% |
| | **NA** | 4% | 12% | 4% | 9% | 7% | 21% | 25% | 8% |

### References

1 for Statistics UI. International standard classification of education: ISCED 2011. *Comp Soc Res* 2012;**30.**:
